# GASTON-Mix: a unified model of spatial gradients and domains using spatial mixture-of-experts

**DOI:** 10.1101/2025.01.31.635955

**Authors:** Uthsav Chitra, Shu Dan, Fenna Krienen, Benjamin J. Raphael

## Abstract

**Motivation:** Gene expression varies across a tissue due to both the organization of the tissue into *spatial domains*, i.e. discrete regions of a tissue with distinct cell type composition, and continuous *spatial gradients* of gene expression within different spatial domains. Spatially resolved transcriptomics (SRT) technologies provide high-throughput measurements of gene expression in a tissue slice, enabling the characterization of spatial gradients and domains. However, existing computational methods for quantifying spatial variation in gene expression either model only spatial domains – and do not account for continuous gradients of expression – or require restrictive geometric assumptions on the spatial domains and spatial gradients that do not hold for many complex tissues.

**Results:** We introduce GASTON-Mix, a machine learning algorithm to identify both spatial domains *and* spatial gradients within each domain from SRT data. GASTON-Mix extends the *mixture-of-experts (MoE)* deep learning framework to a *spatial* MoE model, combining the clustering component of the MoE model with a *neural field* model that learns a separate 1-D coordinate (“isodepth”) within each domain. The spatial MoE is capable of representing any geometric arrangement of spatial domains in a tissue, and the isodepth coordinates define continuous gradients of gene expression within each domain. We show using simulations and real data that GASTON-Mix identifies spatial domains and spatial gradients of gene expression more accurately than existing methods. GASTON-Mix reveals spatial gradients in the striatum and lateral septum that regulate complex social behavior, and GASTON-Mix identifies localized spatial gradients of hypoxia and TNF-*α* signaling in the tumor microenvironment.

## 1 Introduction

Spatially resolved transcriptomics (SRT) technologies measure the gene expression and spatial location of thousands of cells in a 2-D tissue slice. Imaging-based SRT methods, e.g. MERFISH [1] and 10x Xenium [2], measure the expression of hundreds of genes at subcellular resolution, while sequencing-based SRT methods, e.g. Slide-SeqV2 [3], 10x Visium [4], and Seq-Scope [5], measure the expression of genes across the whole transcriptome at near-cellular resolution. The recent development of these high-throughput SRT technologies has enabled the study of the spatial and molecular organization of many biological systems including the brain and tumor microenvironment [6, 7].

Broadly, the major goal of spatial transcriptomic data analysis is to characterize spatial variation in gene expression in a tissue slice from SRT data. Gene expression varies spatially in a tissue due to both the organization of cell types as well as local gradations in cell state. Tissues are organized into *spatial domains*, or discrete regions of a tissue with distinct cell type composition and biological function. For example, the brain is divided into different regions consisting of different types of neurons [8], while tumors often contain multiple spatially coherent clones characterized by transcriptionally distinct subpopulations of cells [9]. At the same time, many genes exhibit continuous *spatial gradients* of expression within a spatial domain. These spatial gene expression gradients may be due to a variety of factors including cell type gradients within a domain, inter-cellular communication, and local changes in cell state and microenvironment, and such gradients may be present even within cells of the same cell type. For instance, several genes exhibit large spatial expression gradients within CA1 pyramidal neurons in the hippocampus [10, 11] and within the striatum region of the brain [12, 13], indicating continuous gradients of neuronal specialization, while in a tumor clone, the expression of genes involved in metabolism or angiogenesis may vary continuously with the distance to the tumor boundary due to oxygen gradients or immune interactions [14].

A large number of computational methods have been introduced for identifying spatial domains from SRT data (e.g. described by [15, 16, 17]). These methods typically assume that (mean) gene expression is *constant* within each domain, and that there are large, discontinuous changes in gene expression across domain boundaries. This assumption is explicitly made by hidden Markov random fields (HMRFs) [18, 19] and is implicitly made by algorithms which cluster cellular embedding vectors, e.g. methods which derive embeddings from graph neural networks (GNNs) [20, 21, 22], variational autoencoders (VAEs) [23], spatial kernels [24], or cell type composition [25]. However, most existing methods for identifying spatial domains do not account for continuous gene expression gradients within a spatial domain; as we demonstrate in this work, not modeling continuous gradients may lead to inaccurate inference of spatial domains.

The standard approach for measuring spatial gradients of gene expression is to derive a 1-D coordinate describing the *relative* position of a spatial location within a spatial domain, and estimate the rate of change of gene expression along this coordinate. Some methods attempt to derive such a 1-D coordinate from low-dimensional embeddings obtained from matrix factorization or deep learning models [26, 27, 28, 29, 30], but because these embeddings do not explicitly model spatial gradients, the learned embeddings do not necessarily correspond to the relative position of a spatial location (e.g. the embeddings may be influenced by non-spatial factors such as cell type or state). Moreover, while there are many methods for identifying spatially variable genes (SVGs; e.g. described by [31]), nearly all of these methods are unable to distinguish spatial gradients from *constant* gene expression within a spatial domain. The few existing approaches for explicitly identifying spatial expression gradients require substantial prior knowledge on the tissue geometry, e.g. manually annotated boundaries [32] or histological tumor annotations [33], in order to estimate the relative position within a given spatial domain. Some of us recently introduced a method, GASTON, that learns a 1-D coordinate called the *isodepth* that continuously varies across the entire tissue slice and can be used to quantify gene expression gradients within spatial domains [34]. The isodepth determines a *“topographic map”* of gene expression, but GASTON assumes that spatial domains are bounded by level sets of the isodepth coordinate. This restrictive geometric assumption often does not hold for complex tissues with an arbitrary arrangement of spatial domains; for example, in this work we derive a tissue and gene expression function which we mathematically prove cannot be modeled by GASTON (Theorem 1). Thus, the problem of fully characterizing spatial variation in gene expression – by both identifying spatial domains *and* a 1-D coordinate within each domain describing spatial gradients of expression – remains unsolved.

We introduce GASTON-Mix, an unsupervised method that simultaneously identifies spatial domains and derives gene expression gradients within spatial domains from SRT data. GASTON-Mix combines the *sparsely-gated, mixture-of-experts (MoE)* deep learning framework [35, 36] with a *neural field* model [37] into a new model that we call a *spatial MoE* model. Sparsely-gated MoE models are widely used in large, transformer-based machine learning models such as ChatGPT due to their large model capacity and efficient training procedure [35], while neural field models are often used in computer graphics and vision for parametrizing physical properties of objects in space, e.g. NeRF [38]. GASTON-Mix uses the gating network of the sparsely-gated MoE model to represent *any* geometric arrangement of spatial domains in a tissue, and GASTON-Mix parametrizes the experts using a neural field model which learns a separate 1-D isodepth coordinate and topographic map for *each* spatial domain.

We show on simulated and real SRT data that GASTON-Mix more accurately identifies spatial domains and spatial gradients of gene expression compared to existing methods. On MERFISH data from the mouse brain, GASTON-Mix reveals spatial gradients in the striatum and lateral septum that regulate complex social behavior. On 10x Genomics Visium data from a breast cancer sample, GASTON-Mix identifies spatial gradients of hypoxia and TNF-*α* signaling that are localized to specific tumor domains.

## 2 Methods

### 2.1 Gene expression functions and spatial gradients

We start by describing a general framework for spatial domains and gene expression gradients in spatially resolved transcriptomics (SRT) data following the exposition by [34]. SRT technologies measure the expression of *G* genes in a tissue slice *T* ⊆ ℝ^2^, which we model using a (vector-valued) *gene expression function* f : *T* →ℝ^*G*^. The vector f (*x, y*) = (*f*_1_ (*x, y*), …, *f*_*G*_ (*x, y*)^⊺^ is the expression of genes *g* = 1, …, *G* at spatial location *x, y* in the tissue slice *T*. The *g*-th component function *f*_*g*_ : ℝ^2^ →ℝ describes the expression of a single gene *g*. For example, a gene *g* whose expression is constant across the tissue slice *T* has a constant expression function *f*_*g*_ (*x, y*) = *c*, while a gene that is differentially expressed in a region *R* ⊆ T might have the expression function *f*_*g*_ (*x, y*) = *c*· 1 _{(*x,y*)∈_*R*_}_ + *c*′· 1 _{(*x,y*) ∉*R*}_.

We model each gene expression function *f*_*g*_ (*x, y*) as a *piecewise, continuously differentiable* function. *Continuously differentiable* functions, or functions *f* whose gradient ∇*f* is continuous, describe continuous gradients of gene expression (e.g. due to gradients in cell type proportion or cellular interactions) while *piecewise* functions allow for discontinuities in expression due to sharp changes in cell type composition or other factors. We assume the expression functions *f*_*g*_ (*x, y*) have the same pieces for all genes *g* = 1, …, *G*, and thus have the form

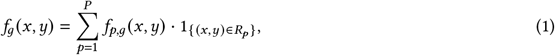

where *R*_1_, …, *R*_*P*_ ⊆ T is a partition of the tissue slice *T* into *P* disjoint regions, which we call *spatial domains*, and *f*_*p,g*_ : *R*_*p*_ → ℝ is a continuously differentiable function describing the expression of gene *g* in domain *R*_*p*_.

A *spatial gradient* describes how gene expression changes across the 2D tissue slice *T*. The spatial gradient for a single gene *g* at spatial location (*x, y*) is given by the gradient ∇*f*_*g*_ (*x, y*) of the expression function *f*_*g*_. More generally, the rows of the Jacobian matrix **J**(**f**)(*x, y*) = [∇*f*_1_ (*x, y*)··· ∇*f*_*G*_ (*x, y*)]^⊺^ ∈ ℝ^*G*×2^ of the gene expression function **f** give the spatial gradient of each gene *g* = 1, …, *G* at spatial location (*x, y*) ∈ *T*. Since the expression function *f*_*g*_ is piecewise continuously differentiable, the spatial gradient ∇*f*_*g*_ is piecewise continuous and may be written is

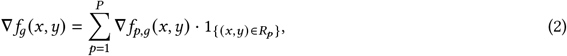

where ∇*f*_*p,g*_ is the gradient of the domain-*p* gene expression function *f*_*p,g*_. We note that ∇*f*_*p,g*_ is a continuous function as *f*_*p,g*_ is continuously differentiable.

### 2.2 GASTON framework and limitations

Directly estimating the spatial gradients ∇*f*_*g*_ for each gene *g* = 1, …, *G* from SRT data is difficult due to the limited spatial resolution and/or limited read depth (e.g. *sparsity*) of current technologies. In our previous work, we proposed to estimate the spatial gradients ∇*f*_*g*_ by making several *global* assumptions on the structure of the spatial gradients ∇*f*_*g*_ and spatial domains *R*_*p*_, implemented in the GASTON algorithm [34]. We briefly describe these assumptions.

First, GASTON assumes that there is a continuous vector field **v** : *T* → ℝ^2^ such that the spatial gradient ∇*f*_*p,g*_ for each gene *g* = 1, …, *G* in spatial domain *p* = 1, …, *P* is proportional to the vector field **v**, i.e.

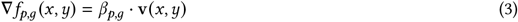

where *β*_*p,g*_ ∈ ℝ is a constant of proportionality and the vector field **v** is called the spatial gradient vector field. In particular, GASTON assumes that the spatial gradient vector field v is a *conservative* vector field, meaning that it has zero curl, i.e. curl(**v**) = 0. Equivalently, the spatial gradient vector field **v** is the gradient of a continuously differentiable scalar function *d* : ℝ^2^ → ℝ, i.e. **v** = ∇*d*. Under these assumptions, the spatial gradients ∇*f*_*p,g*_ are

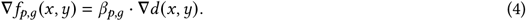

The scalar function *d* (*x, y*) is called the *isodepth* and describes the *“topography”* of the tissue slice *T*, analogous to the elevation in a topographic map of a geographic region.

Second, GASTON assumes that each spatial domain *R*_*p*_ is the union of level sets of the isodepth *d*. Specifically, each domain *R*_*p*_ has the form *R*_*p*_ = {(*x, y*) : *b*_*p*−1_ < *d* (*x, y*) ≤ *b*_*p*_} for some real numbers −∞ = *b*_0_ < *b*_1_ <··· < *b*_p−1_ < *b*_*P*_ = ∞.

GASTON uses these two assumptions to write the gene expression function **f** (*x, y*) as f (*x, y*) = **h** (*d*(*x, y*)), where **h** (*z*) : ℝ → ℝ^*G*^ is a piecewise continuously differentiable function. GASTON approximates the functions *d* (*x, y*) and **h** (*z*) using neural networks. However, there exist biologically realistic expression functions f whose spatial domains *R*_*p*_ *cannot* be described in such a way – and thus cannot be learned by GASTON. We formalize this claim by describing a simple and biologically plausible example gene expression function **f** (*x, y*) which we prove does not satisfy the GASTON assumptions.

#### Gene expression function that cannot be modeled by single isodepth function used in GASTON

Consider a rectangular tissue slice *T* ⊆ ℝ^2^ with *P* ≥ 2 spatial domains where two domains *R*_1_, *R*_2_ are adjacent rectangles. Suppose there exists a gene *g* with domain-1 and domain-2 expression functions *f*_1,*g*_ (*x, y*), *f*_2,*g*_ (*x, y*), respectively, given by

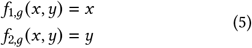

as shown in Figure 1.

**Figure 1:**
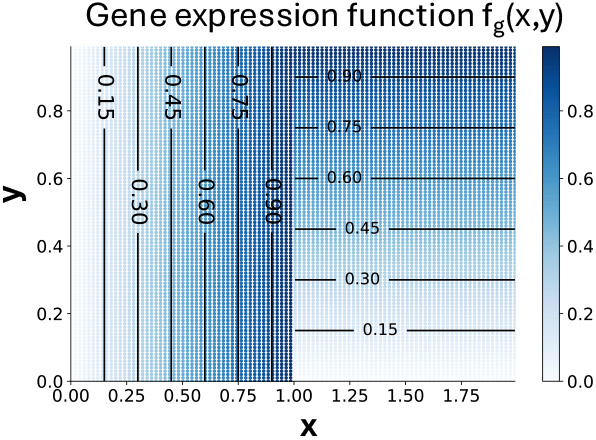
A piecewise continuous gene expression function *f*_*g*_ (*x, y*) which cannot be described with a single, continuous isodepth function *d* (*x, y*) (Theorem 1).

We prove that the two assumptions of GASTON do not hold for this gene expression function **f**. Namely, we show that there does not exist a continuous function *d* : ℝ^2^ → ℝ and piecewise continuous function **h** : ℝ → ℝ^G^ such that **f** (*x, y*) = **h** (*d* (*x, y*)), which is implied by the GASTON assumptions.

##### Theorem 1.

*Let* f : ℝ^2^ → ℝ^*G*^ *satisfy* (5) *for some gene g* ∈ {1, …, *G*}. *There does not exist a continuously differentiable function d* : ℝ^2^ → ℝ *and piecewise continuously differentiable function* **h** : ℝ → ℝ^*G*^ *such that* **f** (*x, y*) = **h**(*d* (*x, y*)).

See the Appendix for a proof. Furthermore, we demonstrate empirically (Section 3.1) that for an expression function f that satisfies (5), GASTON is unable to accurately learn the spatial domains or gene expression gradients.

The fundamental reason that GASTON cannot model such a gene expression function f is because GASTON assumes that the gene expression function **f** has a *global* structure, in that there is a *single*, continuous isodepth function *d* (*x, y*) defined across the entire tissue *T* that describes both the geometry of the spatial domains *R*_*p*_ and the spatial gradients ∇*f*_*p,g*_ within each domain. But as our above example demonstrates, there are scenarios where the gene expression function f does not have such a global structure. These theoretical and empirical observations motivate us to derive a model that relaxes GASTON’s assumption that all spatial domains share a global isodepth function *d* (*x, y*).

### 2.3 Domain-specific spatial topography problem

We derive a model of spatial domains and *local*, domain-specific spatial gradients that relaxes the global assumptions of GASTON described in the previous section.

We start by describing a general model of spatial domains that formalizes the assumptions implicit in many spatial domain identification methods. We define the spatial domains *R*_*p*_ in terms of an *indicator function w*_*p*_ (*x, y*) : ℝ^2^ → {0, 1}, where *w*_*p*_ (*x, y*) = 1 if spatial location (*x, y*) ∈ *R*_*p*_ and *w*_*p*_ (*x, y*) = 0 otherwise. We call *w*_*p*_ (*x, y*) the domain-*p* assignment function. Note that 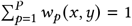 for every spatial location (*x, y*), as the spatial domains *R*_*p*_ are disjoint. The gene expression function *f*_*g*_ may be written as

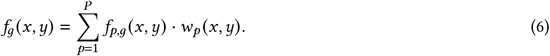

We do not make any global assumptions on the structure of the domain-*p* assignment function *w*_*p*_ *x*(, *y*), as the spatial domains of a tissue may have arbitrary shape and arrangement. In particular, unlike GASTON, we do not assume that the domain-*p* assignment function *w*_*p*_ (*x, y*) is related to the contours of a continuous function *d* (*x, y*). We also relax GASTON’s assumption that there is a global spatial gradient vector field across all domains. Instead, we assume that *each* domain *R*_*p*_ has its own continuous spatial gradient vector field **v**_*p*_ : *R*_*p*_ → ℝ^2^. The spatial gradients 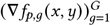 in domain *p* for each gene *g* = 1, …, *G* are proportional to v_*p*_ (*x, y*); i.e., ∇*f*_*p,g*_ (*x, y*) = *β*_*p,g*_· **v**_*p*_ (*x, y*) for some constant *β*_*p,g*_ ∈ ℝ.

We emphasize that we allow *discontinuities* in the domain-specific vector fields **v**_*p*_ on the boundaries *∂R*_*p*_ of spatial domains *R*_*p*_. That is, the tissue-wide spatial gradient vector field 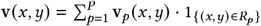 formed by the piecewise sum of the domain-specific vector fields **v**_*p*_ may be discontinuous. In contrast, GASTON’s global model does not allow for discontinuities in the tissue-wide gradient vector field.

Further, we assume that each domain-*p* spatial gradient vector field **v**_*p*_ is a conservative vector field, so that **v**_*p*_ = ∇*d*_*p*_ is the gradient of a continuously differentiable function *d*_*p*_ : *R*_*p*_ → ℝ on the spatial domain *R*_*p*_ which we call the *domain-p isodepth function*. Then the spatial gradients may be written as

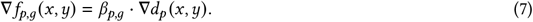

Integrating both sides of (7) yields the following expression for the domain-*p* expression function *f*_*p,g*_:

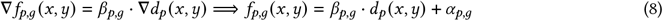

where *α*_*p,g*_, *β*_*p,g*_ ∈ ℝ are constants. If we define the linear function *h*_*p,g*_ (*z*) = *β*_*p,g*_*z* + *α*_*p,g*_, then we may write (8) as

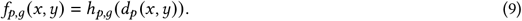

That is, the domain-*p* expression function *f*_*p,g*_ is the composition of the domain-*p* isodepth *d*_*p*_ (*x, y*) and a *linear* function *h*_*p,g*_ (*z*). We call *h*_*p,g*_ (*z*) the domain-*p one-dimensional (1-D)* expression function, as *h*_*p,g*_ (*z*) is a function of a single variable *z* in contrast to the expression function *f*_*p,g*_ (*x, y*) which is a function of two variables *x, y*. Then the gene expression function *f*_*g*_ (*x, y*) across the entire tissue is given by

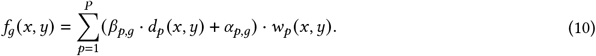

Given SRT data and a number *P* of domains, we compute the maximum likelihood estimators (MLEs) of the spatial domain assignment functions *w*_*p*_ (*x, y*), the domain-*p* isodepth functions *d*_*p*_ (*x, y*), and the domain-*p* 1-D expression functions *h*_*p,g*_ (*z*). The observed SRT data consists of a transcript count matrix A = [*a*_*i,g*_] ∈ ℝ^*N* ×*G*^, where *a*_*i,g*_ is the expression of gene *g* ∈ {1, …, *G*} in spatial location *i* ∈ {1, …, *N*}, and a spatial location matrix **S** = [**s**_*i*_] ∈ ℝ^*N* ×2^, where **s**_*i*_ = (*x*_*i*_, *y*_*i*_) is the *i*-th observed spatial location. We define the Domain-Specific Spatial Topography Problem (DS-STP) as the following maximum likelihood estimation problem.

#### Domain-Specific Spatial Topography Problem (DS-STP)

*Given SRT data* (**A, S**) *and a number P of spatial domains, find indicator functions w*_*p*_ : ℝ^2^ → {0, 1}, *continuously differentiable functions d*_*p*_ : ℝ^2^ → ℝ, *and linear functions h*_*p,g*_ *for domains p* = 1, …, *P and genes g* = 1, …, *G that maximize the log-likelihood of the observed data:*

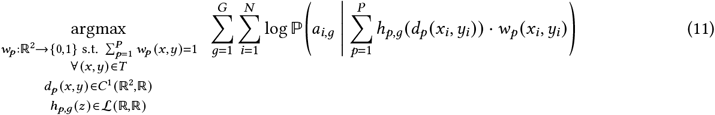

*where C*^1^ (ℝ^2^, ℝ) *is the space of continuously differentiable functions from* ℝ^2^ *to* ℝ *and* ℒ (ℝ, ℝ) *is the space of linear functions from* ℝ *to* ℝ.

The DS-STP problem considerably generalizes the Spatial Topography Problem (STP) from [34]. In the STP, one aims to learn a *single* continuous isodepth function *d* (*x, y*) for all domains, and assumes the spatial domain assignment functions *w*_*p*_ (*x, y*) are level sets of the isodepth function *d*. In contrast, in the DS-STP, we aim to learn *multiple* isodepth functions *d*_*p*_ and do not place any constraints on the spatial domain assignment functions *w*_*p*_ *x*(, *y*).

The DS-STP is a computationally challenging problem to solve as it involves optimizing over spaces of both continuous *and* binary functions. Such an optimization problem is a generalization of *mixed-integer* optimization problems, where one optimizes over continuous and discrete variables (rather than functions). Since mixed-integer optimization problems are NP-hard [39], it is unlikely that one would be able to solve the DS-STP to optimality.

### 2.4 Spatial mixture-of-experts model

We introduce a *spatial mixture-of-experts* model to efficiently and approximately solve the DS-STP. Our new model combines the widely used *sparsely-gated, mixture-of-experts (MoE)* deep learning framework [35, 36] with a neural field model.

Briefly, a sparsely-gated MoE model consists of *P* “expert” neural networks *E*_1_, …, *E*_*P*_; a “gating” (or “routing”) neural network *G* = (*g*_1_, …, *g*_*P*_) ∈ [0, 1]^*P*^; and a fixed integer *k* > 0. Each input s is mapped to a *sparse* linear combination 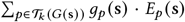, where the linear coefficients *g*_*p*_ (**s**) are given by the gating network *G*(**s**) evaluated on the input **s**, and the sum is taken over the set 𝒯_*k*_ (*G* (**s**)) ⊆ {1, …, *P*} of the *k* largest indices of the vector *G*(**s**); such a sum is sometimes called “top-*k* gating” in the deep learning literature [35]. The expert networks *E*_1_, …, *E*_*P*_ and gating network *G* are trained jointly using backpropagation.

We specifically use a sparsely-gated MoE model where the input s = (*x, y*) to the model is a spatial location (*x, y*) in the tissue slice *T* and the output is the predicted expression vector **a** ∈ ℝ^*G*^ at location (*x, y*). The *p*-th expert *E*_*p*_ : ℝ^2^ → ℝ^*G*^ corresponds to the gene expression function 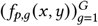 for domain *p*, i.e.

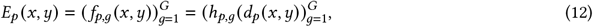

where the second equality uses (8). We parametrize the domain-*p* isodepth function *d*_*p*_ (*x, y*) with a neural network as the universal approximation theorem shows that continuous functions can be closely approximated with neural networks [40]. Since *h*_*p,g*_ (*z*) is a linear function, this is equivalent to parametrizing the *p*-th expert *E*_*p*_ with a neural network whose last hidden layer has a single node and is followed by a linear layer (Figure 2).

**Figure 2:**
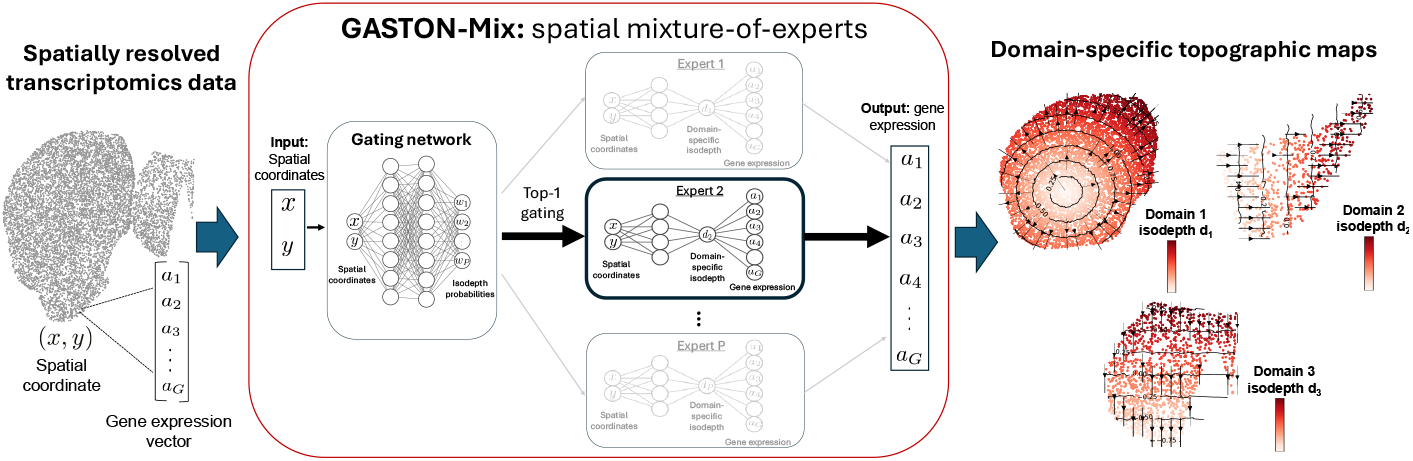
**(Left)** The input to GASTON-Mix is spatially resolved transcriptomics (SRT) data from a 2-D tissue slice, which contains the spatial location (*x, y*) and gene expression a = [*a*_*g*_] ∈ ℝ^*G*^ of thousands of cells in the tissue slice *T*. **(Middle)** GASTON-Mix is an unsupervised, spatial mixture-of-experts (MoE) deep learning model whose input is the spatial coordinate (*x, y*) of a cell and whose output is the predicted gene expression vector a. GASTON-Mix first uses a *gating network G* (*x, y*) = (*w*_1_ (*x, y*), *w*_2_ (*x, y*), …, *w*_*P*_ (*x, y*)) to compute a *soft assignment* of a cell to one of *P* experts, where each expert corresponds to a *spatial domain R*_*p*_ of the tissue slice *T*. GASTON-Mix then uses top-1 gating to make a *hard assignment* of a cell to an expert network *E*_*p*_. Each expert network *E*_*p*_ (*x, y*) predicts gene expression from the spatial coordinate, and has a hidden layer of size 1 corresponding to the *isodepth* coordinate within each domain *R*_*p*_. **(Right)** The domain-specific isodepth coordinate *d*_*p*_ learned by each expert network *E*_*p*_ describes a *topographic map* within each spatial domain *R*_*p*_ and can be used to measure gradients of expression or cell type within the spatial domain.

Further, we use *top-*1 *gating* (i.e., *k* = 1) so that each input spatial location **s** = (*x, y*) is assigned to a single expert *E*_*p*_. Thus each expert ideally corresponds to an individual spatial domain *R*_*p*_ in the tissue. We parametrize the gating function *G* (*x, y*) = (*g*_1_ (*x, y*), …, *g*_*P*_ (*x, y*)) : ℝ^2^ → ℝ^*P*^ with a neural network whose last layer is a softmax, so that the entries 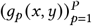 of the gating network are positive and sum to 1, i.e. *g*_*p*_ (*x, y*) > 0 and 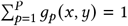. In this way, the gating network *G* (*x, y*) = *g*_1_ (*x, y*), …, *g*_*P*_ (*x, y*): ℝ^2^ → ℝ^*P*^ makes a *soft* assignment of spatial location **s** = (*x, y*) to the *P* spatial domains. Moreover, since we use top-1 gating, the output of our model is the *q* (*x, y*)-th expert *E*_*q*_ (_*x,y*)_ (*x, y*) where *q* (*x, y*) = argmax_*p*=1,…,*P*_ (*g*_1_ (*x, y*), …, *g*_*P*_ (*x, y*)) is the largest index in the entries of the gating network output *G* (*x, y*). That is, the output *E*_*q*(*x,y*)_ (*x, y*) corresponds to a *hard* assignment of each spatial location **s** = (*x, y*) to the *q* (*x, y*)-th spatial domain.

We call our model a *spatial* MoE model as we use neural networks whose inputs are spatial coordinates to parametrize the our expert neural networks *E*_*p*_ (*x, y*) and the gating neural network *G* (*x, y*) of an MoE model. Neural networks whose inputs are spatial coordinates are typically called *neural field* models or *spatial implicit neural representations* in the machine learning literature [37, 41]. To the best of our knowledge, this work and our previous work [34] are the only papers that fit neural field models to spatial transcriptomics data. We note that a recent machine learning paper [42] also proposed a spatial MoE model for weather prediction and forecasting. However, their model differs from ours in two important ways: (1) their gating network does not use spatial information, and so their model does not identify spatial domains and (2) their expert networks do not learn an interpretable 1-D coordinate for measuring spatial gradients.

### 2.5 GASTON-Mix

We implement our spatial MoE model in a package called GASTON-Mix (Figure 2). GASTON-Mix is available at https://github.com/raphael-group/GASTON-Mix with the following implementation choices.

#### Probability model and input features

The MoE model described above can be implemented with different probability distributions for the observed gene expression values *a*_*i,g*_. Following prior work [43, 44], we model the UMI counts *a*_*i,g*_ with a Poisson distribution of the form 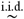 Pois (*U*_*i*_· exp(*f*_*g*_ (*x*_*i*_, *y*_*i*_)) where *U*_*i*_ is the total UMI count at spatial location (*x*_*i*_, *y*_*i*_). In practice, while one could directly solve the DS-STP with all or selected UMI counts, for efficiency we do not directly solve the DS-STP in this way. Instead, we solve the DS-STP using the top generalized linear model principal components (GLM-PCs) [43] under a Gaussian error model. We then use the estimated isodepth functions 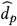 and spatial domain assignment functions 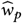 to estimate the 1-D gene expression functions *ĥ*_*p,g*_ for each gene *g* = 1, …, *G* by solving *G*· *P* Poisson regression problems, following [34]. See Appendix for details.

#### Neural network architecture and training

We parametrize the gating network *G* : ℝ^2^ → ℝ^*P*^ with a neural network with two hidden layers of size 20, and we parametrize the domain-*p* isodepth function *d*_*p*_ : ℝ^2^ → ℝ with a neural network with one hidden layer of size 20. We empirically observe that such an architecture is able to represent the arrangement of spatial domains in most tissues as well as the isodepth within each spatial domain without overfitting to the data. We implement GASTON-Mix in PyTorch and we train GASTON-Mix for 50, 000 epochs using a full batch on a single P100 GPU. We use an alternating maximization approach, where we alternate between updating the gating function *G* and the experts *E*_1_, …, *E*_*P*_, as we empirically observe that an alternating optimization approach leads to a lower loss (i.e. larger likelihood (11)). In contrast to GASTON which requires training 30 separate models with different random initializations, we train our spatial MoE model once with a single initialization, with training taking roughly 10 minutes.

#### Initialization and model selection

We often have prior knowledge in the form of approximate spatial domain labels 𝓁_*i*_ ∈ {1, …, *P*} for each spatial location *i* = 1, …, *N*, e.g. from a clustering algorithm such as *k*-means clustering. While these spatial domain labels 𝓁_*i*_ are likely incorrect, it may still be useful to incorporate such prior knowledge into our model and allow the model to “refine” the noisy domain labels during training. In such cases, we first initialize the gating network *G* (*x, y*) by training the gating network *G* to predict the label 𝓁_*i*_ of each spatial location (*x*_*i*_, *y*_*i*_) using a multi-class cross-entropy classification loss [45]. We use the labels from *k*-means clustering [45] in our simulations (Section 3.1) and the labels from CellCharter [23] in our real data evaluation (Section 3.2). We set the number *P* of experts equal to either the ground truth number of spatial domains if it is known (e.g. in the simulations) or the number of clusters determined by CellCharter, as indicated in the results below.

#### Identifying gene expression gradients

A large (absolute) value of the slope *β*_*p,g*_ of the 1-D gene expression function *h*_*p,g*_ (*z*) indicates a continuous gradient in expression for gene *g* in domain *R*_*p*_. Following [34], we say that gene *g* exhibits an *expression gradient in domain R*_*p*_ if the absolute slope |*β*_*p,g*_ | is greater than a threshold *s*_*p*_. We set *s*_p_ to be the fifth percentile of all estimated slope magnitudes 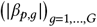 in domain *R*_*p*_.

#### Cell type-specific gradients

Domain-specific gradients in gene expression, i.e. large absolute values of the slope *β*_*p,g*_, may result from spatial variation in cell type proportion. For single-cell data with known cell type annotations, we follow our earlier approach [34] to distinguish between cell type-specific gradients and gradients due to other variation. Briefly, we write the 1-D gene expression function *h*_*p,g*_ (*z*) as 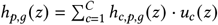 as a sum over cell types *c* = 1, …, *C*, where *u*_*c*_ (*z*) is the proportion of cell type *c* at isodepth *z* and *h*_*c,p,g*_ (*z*) is the *cell type c-specific* 1-D expression function. Given cell type annotations 𝓁_*i,c*_ ∈ {0, 1}, where 𝓁_*i,c*_ = 1 if spatial location *i* contains cell type *c* and 𝓁_*i,c*_ = 0 otherwise, we learn a (linear) cell type-specific 1-D expression function *h*_*c,p,g*_ (*z*) = *β*_*c,p,g*_· *d*_*p*_ (*x, y*) + *α*_*c,p,g*_ for each cell type *c*, across all genes *g* with an expression gradient in domain *R*_*p*_ as defined above.

A large absolute value of the *cell type-specific* slope *β*_*c,p,g*_ for any cell type *c* describes an expression gradient in gene *g* expression in domain *R*_*p*_ that is specific to cell type *c*, and thus is not driven by a gradient in cell type *c* proportion, while a small value of the slope *β*_*c,p,g*_ for all cell types *c* indicates that the expression gradient is driven by a gradient in cell type proportion. See [34] for specific details.

## 3 Results

### 3.1 Evaluation on simulated SRT data

We first evaluated GASTON-Mix on simulated SRT data. We compared GASTON-Mix to four recent methods for analyzing SRT data: BANKSY [24], CellCharter [23], GraphST [22], and GASTON [34]. BANKSY, CellCharter, and GraphST are methods for spatial domain identification which do not explicitly model gene expression gradients within domains, while GASTON is primarily designed for identifying spatial gradients and makes restrictive global assumptions on tissue geometry.

We simulated SRT data S, A on a rectangular tissue *T* consisting of *N* = 10, 000 spatial locations s_*i*_ = (*x*_*i*_, *y*_*i*_) arranged in a 100 ×100 grid. The tissue*T* consists of *P* = 2 spatial domains *R*_1_, *R*_2_ arranged in a “checkerboard” pattern where each domain *R*_*i*_ consists of alternating squares (Figure 3A). This geometric arrangement of spatial domains matches the simulation set-up from [25] which found that such a checkerboard arrangement was difficult for many existing SRT methods – including GASTON and BANKSY – to model. The domain-1 isodepth function *d*_1_ (*x, y*) is the distance from the nearest left edge of a square in domain 1, and the domain-2 isodepth function *d*_2_ (*x, y*) is the distance from the nearest bottom edge of a square in domain 2 (Figure 3B).

**Figure 3:**
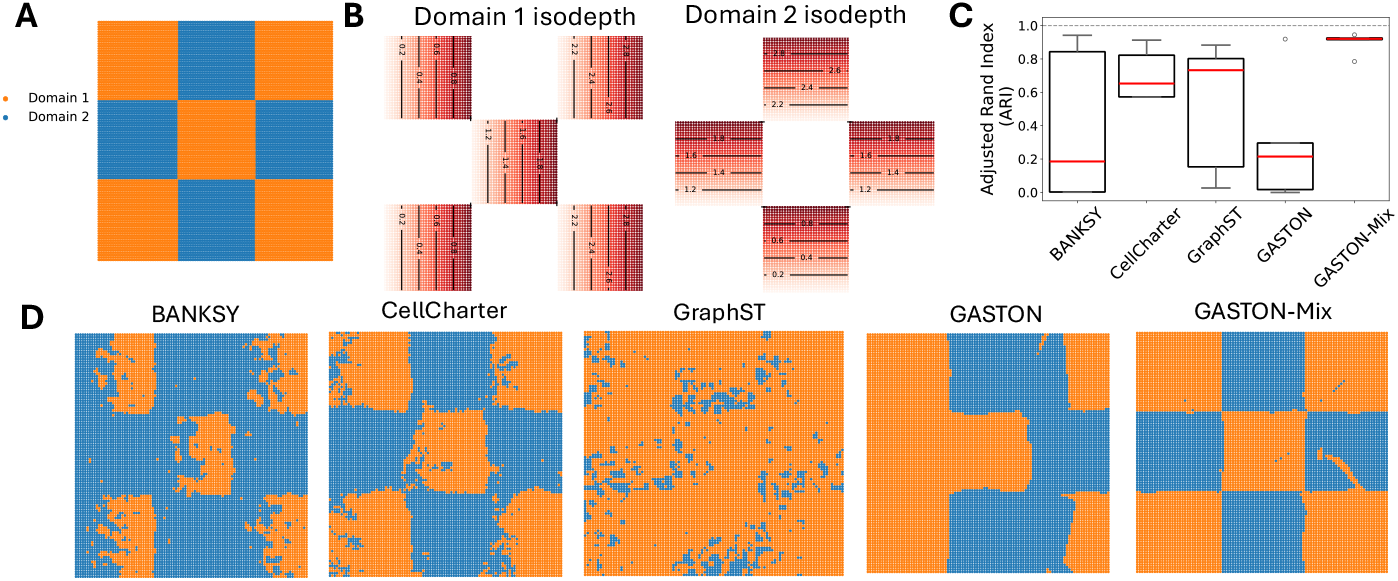
**(A)** Spatial domains of simulated SRT dataset in a rectangular tissue *T* = *R*_1_ ∪ *R*_2_ with two domains *R*_1_ (orange) and *R*_2_ (blue) arranged in a “checkerboard” pattern. **(B)** Domain-specific isodepth functions *d*_1_ (*x, y*) and *d*_2_ (*x, y*) for domains 1 (left) and 2 (right) respectively. Lines denote contours of constant isodepth. **(C)** Adjusted Rand index (ARI) of spatial domains identified by BANKSY [24], CellCharter [23], GraphST [22], GASTON [34], and GASTON-Mix over five simulated instances. Red lines indicate median ARI. **(D)** Spatial domains identified by each method for a single simulated instance.

We simulate the expression values 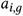 of *G* = 3*G*^′^ genes using three different classes of expression functions *f*_*g*_ (*x, y*) following (10) that satisfy the assumptions of Theorem 1. The first class consists of linear functions of the domain-1 isodepth *d*_1_ (*x, y*); the second class consists of linear functions of the domain-2 isodepth *d*_1_ (*x, y*); and the third class consists of linear functions of both the domain-1 isodepth *d*_1_ (*x, y*) and the domain-2 isodepth *d*_2_ (*x, y*) (Figure S1A, Appendix). We draw gene expression values 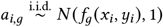 from a normal distribution centered at *f*_g_ (*x, y*) for *G*^′^ = 5 genes from each class (Figure S1B). When running all methods, we omit the pre-processing steps (i.e. PCA or GLM-PCA) as the gene expression values are already low-dimensional and normally distributed, allowing us to solely compare the different algorithms.

We find that GASTON-Mix achieves much higher accuracy in spatial domain identification (mean ARI ≈ 0.85), measured in terms of the adjusted Rand index (ARI), compared to BANKSY, CellCharter, GraphST, or GASTON (mean ARI ≈ 0.37, 0.69, 0.50, 0.40, respectively; Figure 3C). The lower ARI of BANKSY, GraphST, and CellCharter is likely because these methods do not explicitly model gene expression gradients, while GASTON’s poor performance is because the simulated gene expression functions *f*_*g*_ (*x, y*) and spatial domains *R*_*i*_ do not satisfy GASTON’s global isodepth assumption (Theorem 1). We note that while BANKSY claims to model *“gradients in gene expression in [cellular] neighborhoods”* [24], the BANKSY domains have the lowest ARI which suggests that BANKSY is not accurately modeling gradients. In contrast, GASTON-Mix’s local model of tissue geometry and expression gradients allows for accurate inference of spatial domains.

We next evaluated the ability of GASTON-Mix and other methods to learn spatial gradients. We specifically evaluated how well each method learns the domain-specific isodepth functions *d*_*i*_ (*x, y*), which provide coordinates for quantifying spatial gradients. We measure the mean Kendall’s *τ* correlation between the true domain-specific isodepth coordinates *d*_1_ (*x, y*), *d*_2_ (*x, y*), and the isodepth(s) learned by GASTON and GASTON-Mix. While GraphST does not explicitly learn a coordinate for measuring gene expression gradients, GraphST learns a 20-dimensional embedding vector at each spatial location, and we evaluate GraphST using the largest Kendall’s *τ* correlation between each component of the GraphST embedding and the domain-specific isodepth coordinates *d*_1_ *x*(, *y*), *d*_2_ (*x, y*). We do not compare against BANKSY and CellCharter which only identify spatial domains and do not learn a coordinate within each domain for measuring gradients.

We find (Figure 4A) that GASTON-Mix has larger Kendall’s *τ* correlation (*τ* ≈ 0.33) with the true domain-specific isodepths than GASTON (*τ* ≈ 0.27) or GraphST (*τ* ≈ 0.17). GraphST is unable to learn a spatially smooth embedding (Figure 4B) while the GASTON isodepth function is constant in several of the squares (Figure 4C), likely because there does not exist a continuous isodepth function *d* that describes the describes the gene expression function **f**(*x*, *y*) (see Theorem 1). On the other hand, the GASTON-Mix domain-specific isodepths smoothly vary across the domains (Figure 4D), showing the advantages of learning *local* topographic maps.

**Figure 4:**
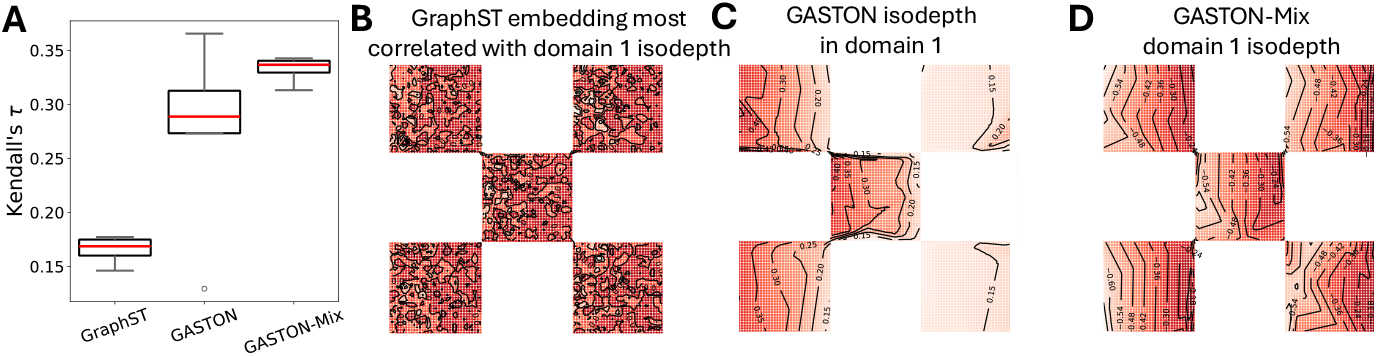
**(A)** Kendall’s *τ* coefficient between simulated domain-specific isodepth (shown in 3B) and the estimated isodepth from GraphST, GASTON, and GASTON-Mix. **(B-D)** (B) GraphST embedding most correlated with the true isodepth, (C) GASTON isodepth, and (D) GASTON-Mix domain-1 isodepth in the true domain 1. Curves denote contours of constant value.

### 3.2 Analysis of mammalian forebrain structures

We next evaluated GASTON-Mix on a sample of anterior structures of the mouse forebrain, where MERFISH was applied to measure the expression of 1, 122 genes in 9, 696 cells [6]. This data is highly sparse, with a median of 312 measured transcripts per cell. The mouse forebrain has a complex geometry, consisting of several spatial domains of differing sizes and relative positions including the striatum (consisting of the dorsal caudoputamen and ventrally located nucleus accumbens), olfactory tubercle, and lateral septum, as labeled in the Allen Mouse Brain Common Coordinate Framework (CCF, Figure 5A) [46]. Furthermore, earlier prominent biological studies have reported several gradients of gene expression, neurochemicals, and connectivity across different regions of the striatum and lateral septum and in different directions, including the dorsolateral-ventromedial gradient in the striatum [47, 48] and the dorsal-ventral gradient in the lateral septum [49] (Figure 5A), which cannot be identified from non-spatial data [12, 13]. Thus, we hypothesized that this MERFISH sample would pose a challenge for existing spatial domain identification methods that do not account for such gradients, but also for algorithms like GASTON which make global, geometric assumptions about tissue geometry.

**Figure 5:**
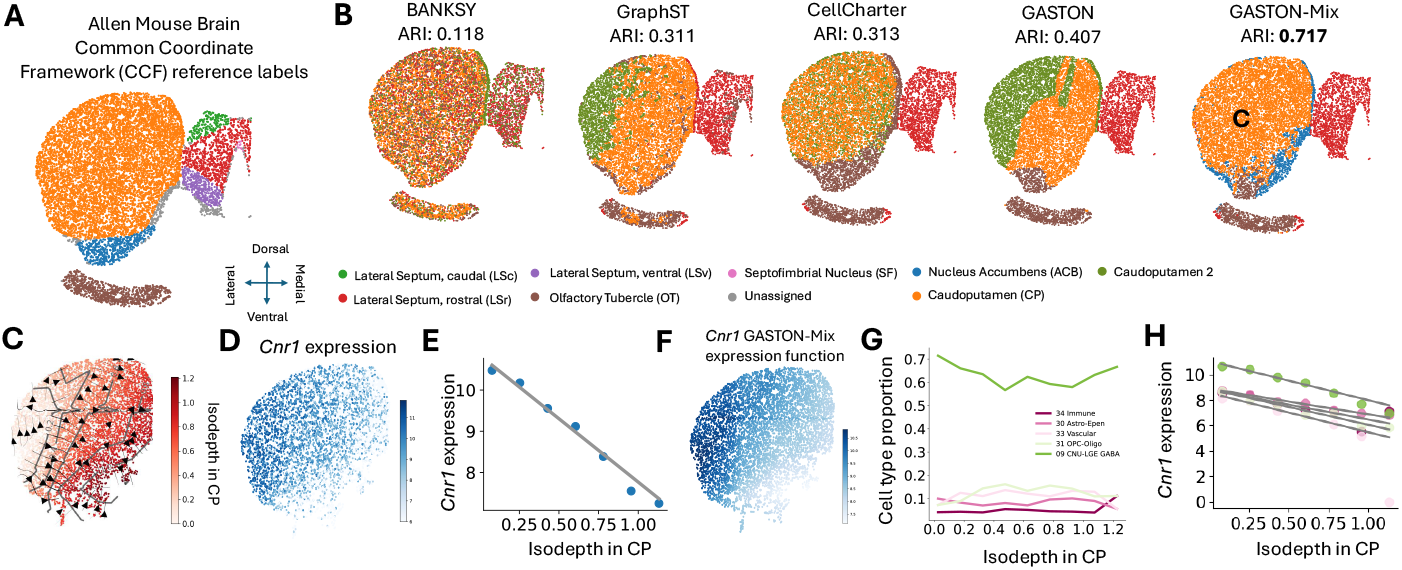
**(A)** Anatomical labels of each cell in a MERFISH mouse anterior forebrain sample from the Allen Mouse Brain Common Coordinate Framework (CCF) [6]. **(B)** Spatial domains and adjusted Rand index (ARI) compared to CCF labels in (A) for five different methods: BANKSY [24], GraphST [22], CellCharter [23], GASTON [34], and GASTON-Mix. **(C)** GASTON-Mix isodepth 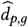 for GASTON-Mix-inferred caudoputamen (CP, labeled orange region in (B)). **(D)** *Cnr1* expression shown in log counts per million (CPM). **(E)** *Cnr1* expression versus CP-specific isodepth 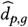 learned by GASTON-Mix. **(F)** GASTON-Mix *Cnr1* expression function 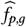 in CP domain. **(G)** Cell type proportion versus CP-specific isodepth 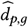 for five most prevalent cell types. **(H)** GASTON-Mix cell type-specific *Cnr1* expression functions ĥ_*c,p,g*_ colored by cell types in (G).

We compared the spatial domains identified by GASTON-Mix to the domains identified by BANKSY, CellCharter, GraphST, and GASTON. We find that GASTON-Mix has larger agreement with the CCF domain labels, measured by the adjusted Rand index (ARI, Figure 5B), compared to the four other methods (GASTON-Mix ARI = 0.717 versus ARI < 0.41 for other methods). In particular, GASTON-Mix is the only method which resolves the caudoputamen (CP, orange domain in Figure 5A) as a single domain while other methods split this domain into two or more domains, with the Banksy and CellCharter output showing little spatial coherence in the orange domain. The reason the other methods fail to identify the CP domain is likely because of the prominent dorsolateral-ventromedial gradient in the domain [47, 48, 12]. We also note that GASTON, a method primarily developed to identify spatial gradients, has a larger ARI than the other methods which are specialized for spatial domain identification (GASTON ARI = 0.407 versus BANKSY, GraphST, CellCharter ARI < 0.32, Figure 5B), demonstrating the difficulty that existing domain identification methods have when the data contains large gene expression gradients, e.g. the dorsolateral-ventromedial gradient in the CP domain.

The domain-specific isodepth functions 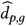 learned by GASTON-Mix reveal several biologically relevant expression gradients that are not readily apparent from the raw MERFISH data. In particular, GASTON-Mix learns a domain-specific isodepth *d*_*p,g*_ that smoothly varies across the domain in the GASTON-Mix-inferred caudoputamen (CP, Figure 5C), allowing for the identification of genes *g* with CP-specific expression gradients, i.e. with large absolute slopes |*β*_*p,g*_|. Such a coordinate is not learned by SRT embedding methods which learn spatially incoherent embeddings, e.g. the GraphST embeddings in the CP domain (Figure S2). GASTON-Mix identifies *Cnr1* – a reported marker for the dorsolateral-ventromedial striatum gradient [50, 51] – as having a CP-specific expression gradient (Methods). *Cnr1* has sparse expression in the domain with a median expression of 1 per cell (Figure 5D). GASTON-Mix aggregates expression across contours of constant isodepth (Figure 5C) and learns a 1-D linear gene expression function *h*_*p,g*_ *z* (Figure 5E) and a 2-D expression function *f*_*p,g*_ (*x, y*) = *h*_*p,g*_ (*d*_*p*_ (*x, y*)) (Figure 5J) that more readily displays the continuous expression gradients compared to the sparse expression. We also find that there are no large gradients of cell type proportion in the CP (Figure 5G; cell type annotations from [6]) and that *Cnr1* exhibits a *cell type-specific* expression gradient across all cell types in the CP domain (Methods). This suggests that the *Cnr1* expression gradient is not driven by variation in cell type proportion and is likely due to other biological causes, e.g. gradients in excitatory input [47]. We also observe similar patterns for other candidate markers for the dorsolateral-ventromedial striatum gradient in different species (Figure S3).

GASTON-Mix also identifies biologically relevant spatial gradients in the lateral septum domain; such gradients have previously been observed in other spatial imaging data and are hypothesized to be involved in complex social behaviors [52, 53, 49]. See Appendix, Figure S4 for details.

Overall, our results demonstrate that while existing SRT algorithms are challenged by the presence of spatial expression gradients, GASTON-Mix’s spatial MoE framework and local isodepth model accurately identifies spatial domains and reveals biologically meaningful spatial gradients of gene expression.

### 3.3 Gradients in breast cancer tumor

We applied GASTON-Mix to a 10x Genomics Visium SRT sample of a human breast cancer (invasive ductal carcinoma) where the expression of 36, 601 genes was measured in 3, 798 spots [54]. Physical and biochemical gradients within a tumor have been previously shown to promote cancer progression and metastasis [55]. GASTON – the current state-of-the-art method for identifying spatial gradients from SRT data – requires the spatial domains of a tissue to satisfy restrictive geometric assumptions [34]. However, the pathologist annotations for this cancer sample (Figure 6A) suggest that its spatial domains have a complex geometry that does not satisfy the GASTON assumptions. Thus, we hypothesized that the more flexible model of GASTON-Mix would allow us to identify spatial domains and gradients inside this tissue.

**Figure 6:**
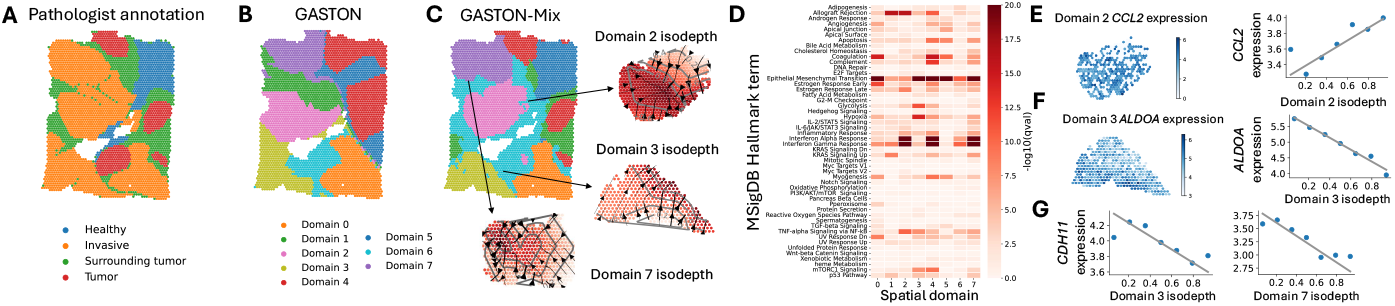
**(A)** Pathologist annotation [54] for each spot in a 10x Genomics Visium breast cancer sample. **(B)** Spatial domains identified by GASTON [34]. **(C)** Spatial domains identified by GASTON-Mix and inferred domain-specific isodepth functions *d*_*p*_ for domains *p* = 2, 3, 7. **(D)** Enrichment for hallmark cancer gene sets reported by gene set enrichment analysis (GSEA) [59] for genes with large expression gradients in each of the *P* = 8 domains identified by GASTON-Mix. **(E-F)** (Left) Expression and (Right) Isodepth versus expression for (E) *CCL2* in domain 2 and (F) *ALDOA* in domain 3. **(G)** *CDH11* expression versus (Left) domain 3 isodepth and (Right) domain 7 isodepth.

We first compared the *P* = 8 domains identified by GASTON and GASTON-Mix. We observe that the GASTON spatial domains (Figure 6B) have noticeably less agreement with the pathologist annotations than the GASTON-Mix domains (Figure 6C), both visually and quantitatively (GASTON ARI ≈ 0.10 versus GASTON-Mix ARI ≈ 0.19 compared to pathologist annotations). This provides evidence that this data does not satisfy the GASTON assumptions, and demonstrates the improved flexibility of GASTON-Mix in modeling arbitrary arrangements of spatial domains compared to GASTON.

We next used GASTON-Mix to identify *shared* and *domain-specific* spatial gradients across the different domains of the cancer sample. Specifically, we identified the set of genes with domain *p*-specific expression gradients across domains *p* = 1, …, *P* (Methods) and subsequently examined the enrichment of these genes in known cancer hallmark gene sets (Figure 6D). We observe that some hallmark sets are enriched across many domains, i.e. they are *shared* across domains, while other sets are enriched in specific domains, i.e. they are *domain-specific*. For example, genes involved in TNF-*α* signaling such as *CCL2* [56] exhibit spatial gradients specifically in invasive domain *R*_2_ (Figure 6E) and genes involved in hypoxia such as the hypoxia-inducible gene *ALDOA* [57] exhibit spatial gradients specifically in invasive domain *R*_3_ (Figure 6F). On the other hand, gradients of genes involved in the epithelial-mesenchymal transition (EMT), such as *CDH11* [58], are present across multiple domains including invasive domains *R*_3_ and *R*_7_ (Figure 6G). We note that the spatial gradients identified by GASTON-Mix would be challenging to identify using existing SRT methods which do not learn coordinates that smoothly vary within domains (e.g. GraphST embeddings in Figure S5).

In this way, GASTON-Mix reveals spatial gradient patterns that characterize the different tumors of a cancer sample.

## 4 Discussion

We introduce GASTON-Mix, a deep learning algorithm for jointly learning spatial domains and gradients of gene expression within each spatial domain from SRT data. We derive a novel *spatial mixture-of-experts (MoE)* model which combines spatial clustering from the standard MoE model with a spatial neural field model that learns a 1-D coordinate – the *isodepth* – within each spatial domain and gives the gradient direction of the expression of individual genes. Our spatial MoE model is capable of representing arbitrary geometric arrangements of spatial domains in a tissue, while the isodepth coordinates within each spatial domain describe *domain-specific topographic maps* that generalize the global topographic maps learned by our previous method GASTON [34]. We show that by modeling gene expression gradients, GASTON-Mix more accurately infers spatial domains compared to existing methods. Moreover, by learning local isodepth coordinates for each spatial domain, GASTON-Mix overcomes the sparsity of existing SRT technologies and reveals domain-specific spatial gradients.

We anticipate many future directions for our work. These include *statistical testing* for whether a domain contains any gene expression gradients, i.e. testing whether the domain-specific isodepth is *constant*; further studying the biological causes of the gene expression gradients identified by GASTON-Mix, e.g. by identifying cell-cell interactions that mediate expression gradients [60, 61]; and relaxing the GASTON-Mix assumptions to allow for inference of *multiple* gradient directions within a spatial domain. Ultimately, we believe that GASTON-Mix’s unified model of spatial domains and gradients provides a general framework for characterizing spatial variation in gene expression across complex tissues.

## Appendix

### A Proof of Theorem 1

Without loss of generality, we assume the domains *R*_1_ and *R*_2_ are squares of side-length 1 that are side-by-side, where the bottom-left corner of square *R*_1_ is 0, 0 and the bottom-left corner of square *R*_2_ is (1, 0), as depicted in Figure 1. We first prove the following lemma.

#### Lemma 1.

*Let* **f** : ℝ^2^ → ℝ^*G*^ *be a function such that* **f** (*x, y*) = h(*d* (*x, y*)) *for continuous function d* : ℝ^2^ → ℝ *and piecewise continuous function* **h** : ℝ → ℝ^*G*^, *and suppose there exists g* ∈ {1, …, *G*} *such that f*_*g*_ *has the form in* (5). *Then for any x*_0_ ∈ [0, 1], *the function e* (*y*) = *d* (*x*_0_, *y*) : [0, 1] → ℝ *is constant. Similarly, for any y*_0_ ∈ [1, 2], *the function e*^′^(*x*) = *d* (*x, y*_0_) *is constant*.

*Proof*. The second statement follows from a similar argument as the first statement, so we prove the first statement. Assume for the sake of contradiction that the first statement is not true. Then there exists *x*_0_ ∈ [0, 1] and 0 ≤ *y* < *y*^′^ ≤ 1 such that *d* (*x*_0_, *y*) ≠ *d* (*x*_0_, *y*^′^). Without loss of generality we may assume that *x*_0_ = 0, and moreover that *d* (0, *y*) < *d* (0, *y*^′^) (e.g. by negating *d*).

Let 0 < *ε* < *d* (0, *y*^′^) −*d* (0, *y*) be sufficiently small. By Lemma 1, we have that *d* (0, *y*) ≠ *d* (*x* ^′^, *y*) for any *x*^′^> 0. By continuity of *d* in *R*_1_, there exists an *x* ^′^ > 0 such that *d* (*x* ^′^, *y*) = *d* (0, *y*) + *ε*. Furthermore, by the intermediate value theorem for the univariate function *y* ↦→ *d* (0, *y*), there exists a *y*^′′^ ∈ (*y, y*^′^) such that *d* (0, *y*^′′^) = *d* (0, *y*) + *ε*.

Thus, we have that *d* (*x* ^′^, *y*) = *d* (0, *y*^′′^) = *d* (0, *y*) + *ε*, which implies

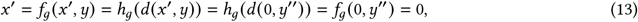

where the first and last equalities follow by definition of *f*_*g*_, the second and second-to-last equalities follow by definition of **f**, and the third equality follows by definition of *x* ^′^ and *y*^′′^. However this contradicts our assumption that *x* ^′^ > 0.

We now proceed to the proof of the Theorem.

*Proof of Theorem 1*. Assume for the sake of contradiction that there exists a continuous function *d* : ℝ^2^ → ℝ and piecewise continuous function h : ℝ → ℝ^*G*^ such that f : ℝ^2^ → ℝ^*G*^ can be written as f (*x, y*) = h(*d* (*x, y*)) for all (*x, y*) ∈ *T*.

From Lemma 1, we have that *d* (*x, y*) is a function of the *x*-coordinate in *R*_1_ and is a function of the *y*-coordinate in *R*_2_. Thus, we may write 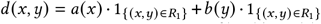 for continuous functions *a*(*x*), *b* (*y*) : ℝ → ℝ.

Now, on the line *x* = 1 where the squares overlap, then *d* (1, *y*) | = *a*(1). Thus, *d* (1, *y*) is constant for all *y* ∈ [0, 1]. But by Lemma 1, for any 0 ≤ *y* < *y*^′^ ≤ 1, we have that *d* (1 + *ε, y*) ≠ *d* (1 + *ε, y*^′^); taking the limit as *ε* → 0 implies that *d* (1, *y*) ≠ *d* (1, *y*^′^) which is a contradiction.

### B GASTON-Mix implementation

We solve the DS-STP using the top generalized linear model principal components (GLM-PCs) [43] under a Gaussian error model. We then use the estimated isodepth functions 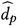 and spatial domain assignment functions 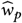 to estimat the 1-D gene expression functions ĥ_*p,g*_ – and thus estimate the spatial domains 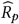– for each gene *g* = 1, …, *G* by solving *G · P* Poisson regression problems, following the procedure of [34]. Our use of GLM-PCA is justified by the following observation from [32]: for SRT data (**A, S**) where the UMI counts *a*_*i,g*_ follow a Poisson distribution with mean *f*_*g*_ (*x*_*i*_, *y*_*i*_) defined by (10), then the top-2*P* GLM-PCs are approximately piecewise linear with Gaussian noise.

Specifically, given SRT data (A, S), we first compute the top-2*P* GLM-PCs u_*j*_ = [*u*_*i,j*_] ∈ℝ^*N*^ for *j* = 1, …, 2*P*. We then use the algorithm described in Section 2.4 to compute the following MLE under the Gaussian error model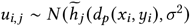 on the GLM-PCs, where 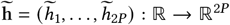 a 1-D GLM-PC function. That is, we solve

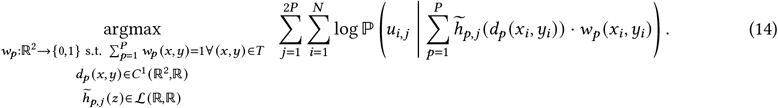

Solving (14) yields estimated isodepth functions 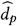 and spatial domain assignment functions 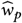. We then estimate the 1-D gene expression functions *h*_*p,g*_ for each gene *g* = 1, …, *G* by solving (14) fixing the isodepth functions 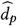 and spatial domain assignment functions 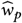, i.e.

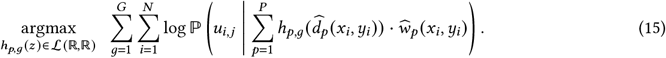

We solve (15) using Poisson regression for each individual gene *g* = 1, …, *G* and spatial domain *p* = 1, …, *P*. This is equivalent to solving *GP* Poisson regression problems. See [32, 34] for details.

### C Simulation set-up

We simulate the expression values *a*_*i,g*_ of *G* = 3*G*^′^ genes using three different classes of expression functions *f*_*g*_ (*x, y*) of the form (10): (1) a linear function *f*_*g*_ (*x, y*) = *α*_1,*g*_*d*_1_ (*x, y*) + *β*_1,*g*_ of only the domain-1 isodepth function *d*_1_ (*x, y*); (2) a linear function *f*_*g*_ (*x, y*) = *α*_2,*g*_*d*_2_ (*x, y*) + *β*_2,*g*_ of only the domain −2 isodepth function *d*_2_ (*x, y*); and (3) a linear function *f*_*g*_ (*x, y*) = *α*_1,*g*_*d*_1_ (*x, y*) + *β*_1,*g*·_ *w*_1_ (x, *y*) + *α*_2,*g*_*d*_2_ (*x, y*) + *β*_2,*g*·_ *w*_2_ (x, *y*) of both domain-specific isodepths *d*_1_ (*x, y*), *d*_2_ (*x, y*). We draw the gene expression function parameters *α*_*p,g*_, *β*_*p,g*_ ~ *U* (−1, 1) uniformly at random for each gene *g* = 1, …, *G* and domains *p* = 1, 2 (Figure S1A). We draw gene expression values 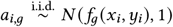 from a normal distribution centered at *f*_*g*_ (*x, y*) for *G*^′^ = 5 genes from each class (Figure S1B).

### D Spatial gradients in lateral septum

GASTON-Mix identifies biologically relevant spatial gradients in the lateral septum (LS) domain (Figure 5A). Spatial gradients in the LS have previously been observed in other spatial imaging data and are hypothesized to be involved in complex social behaviors including reward processing and motor learning [52, 53, 49]. In particular, GASTON-Mix identifies two genes which were recently observed to be markers of several subtypes of lateral septum neurons in a separate MERFISH lateral septum dataset [49], *Ano2* and *Lamp5*, as having large LS-specific gradients (Figure S4C-J). These gradients may be driven by gradients in cell type proportion, e.g. the decreasing gradient in “LSX GABA” cell type proportion (Figure S4B).

**Figure S1:**
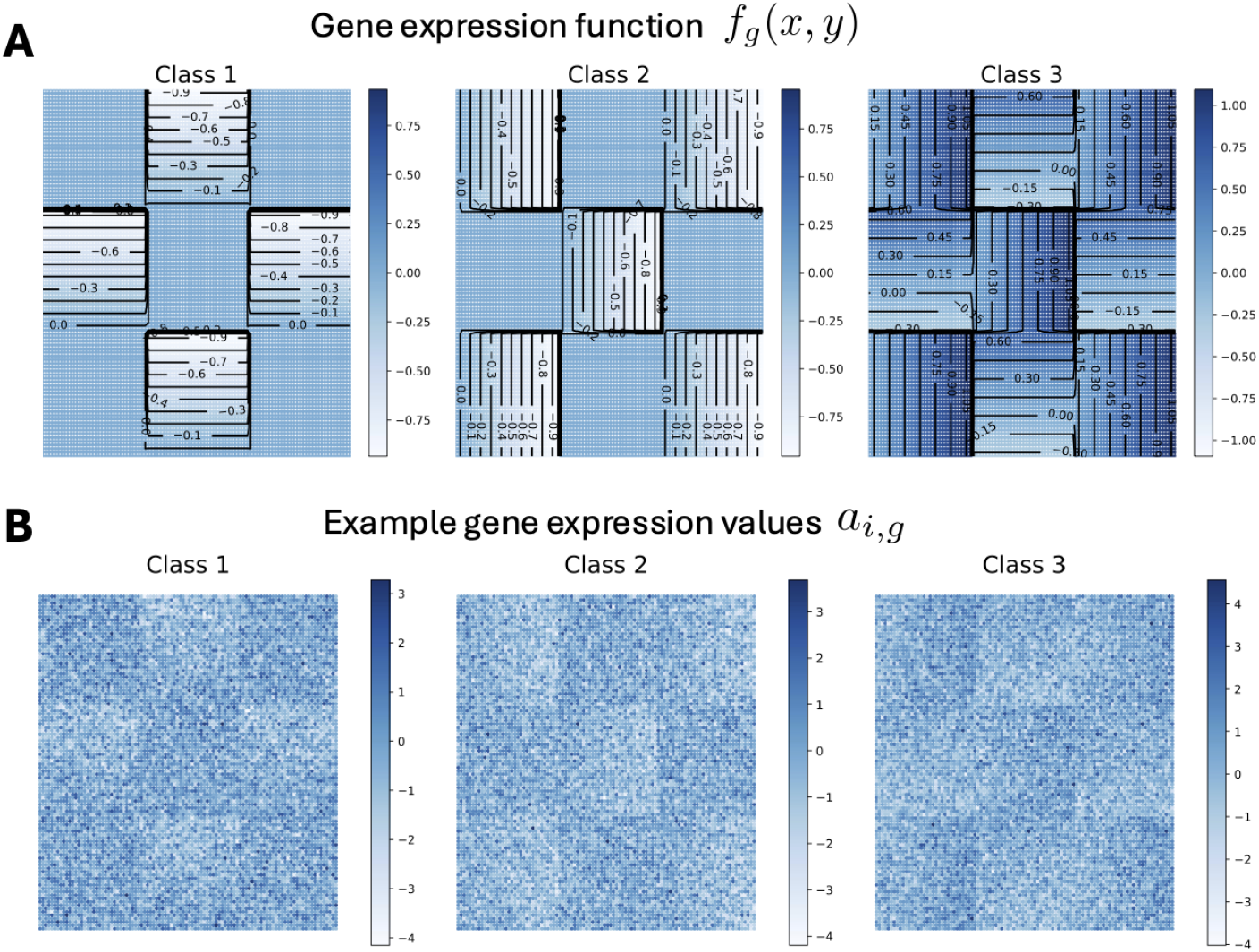
**(A)** Three classes of gene expression functions *f*_*g*_ (*x, y*). **(B)** Three examples of simulated gene expression *a*_*i,g*_ from each of the classes of gene expression functions shown in (A). **(C)** GraphST embeddings most correlated with domain 1 isodepth *d*_1_ (*x, y*) (left) and domain 2 isodepth *d*_2_ (*x, y*) (right).

**Figure S2:**
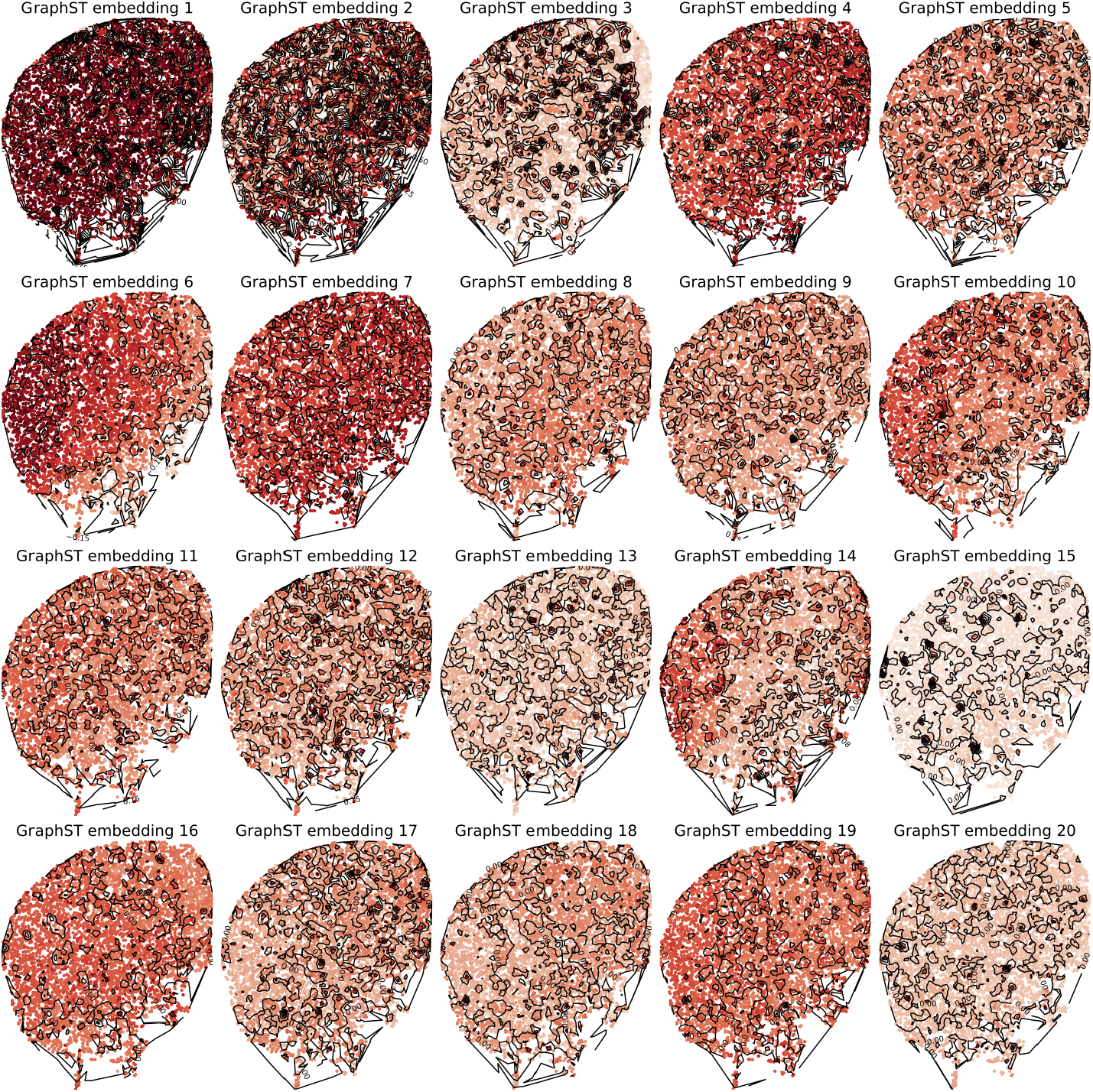
GraphST embedding components for GASTON-Mix-identified CP domain in MERFISH mouse forebrain. Curves connect spots of equal embedding value.

**Figure S3:**
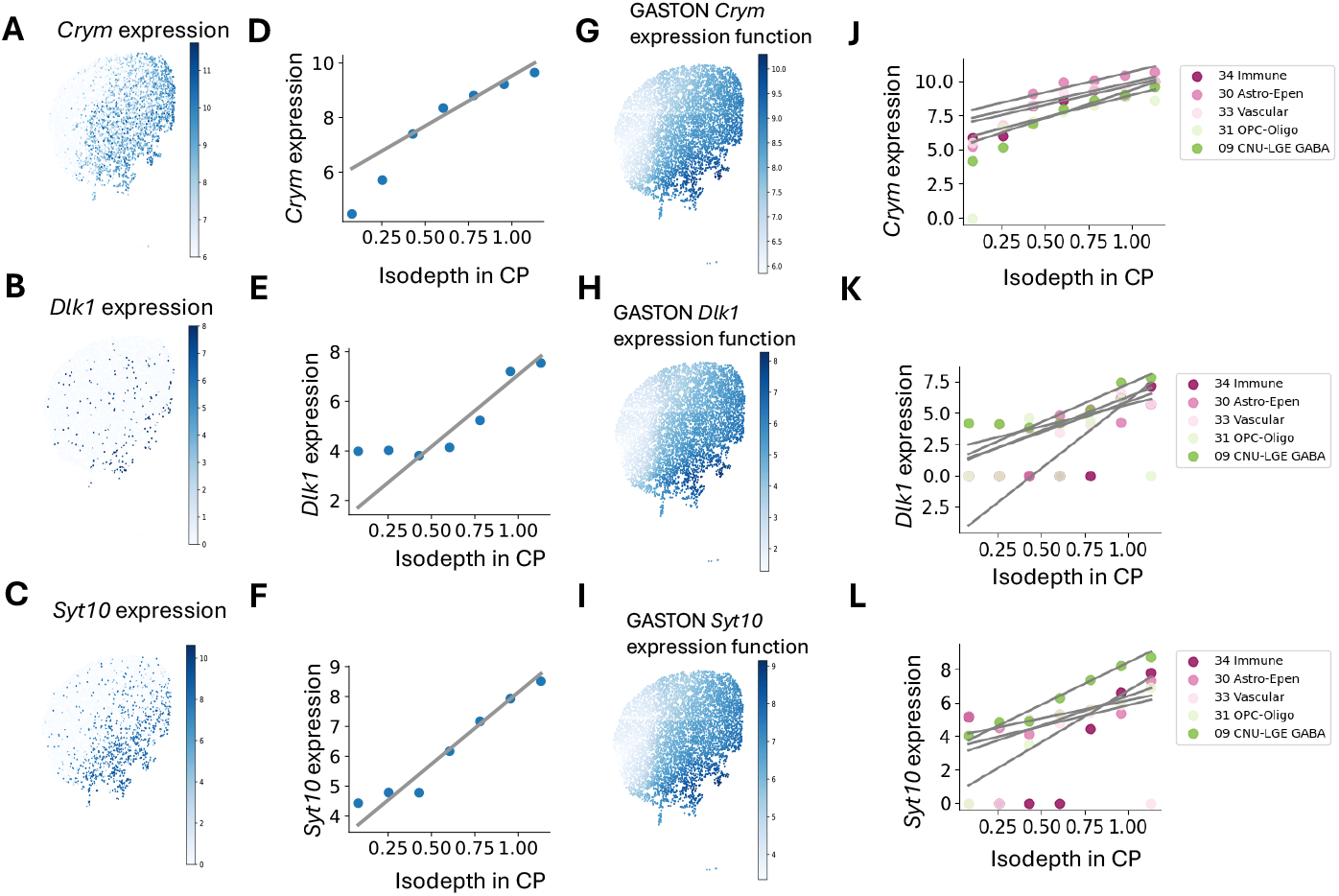
**(A-C)** Gene expression in CP domain for (A) *Crym*, (B) *Dlk1*, and (C) *Syt10*. **(D-F)** Isodepth in CP versus expression for (D) *Crym*, (E) *Dlk1*, and (F) *Syt10*, three candidate markers for the dorsolateral-ventromedial striatum gradient [51]. **(G-I)** GASTON CP-specific expression function for (G) *Crym*, (H) *Dlk1*, and (I) *Syt10*. **(J-L)** Isodepth in CP versus cell type-specific expression functions for (J) *Crym*, (K) *Dlk1*, and (L) *Syt10*.

**Figure S4:**
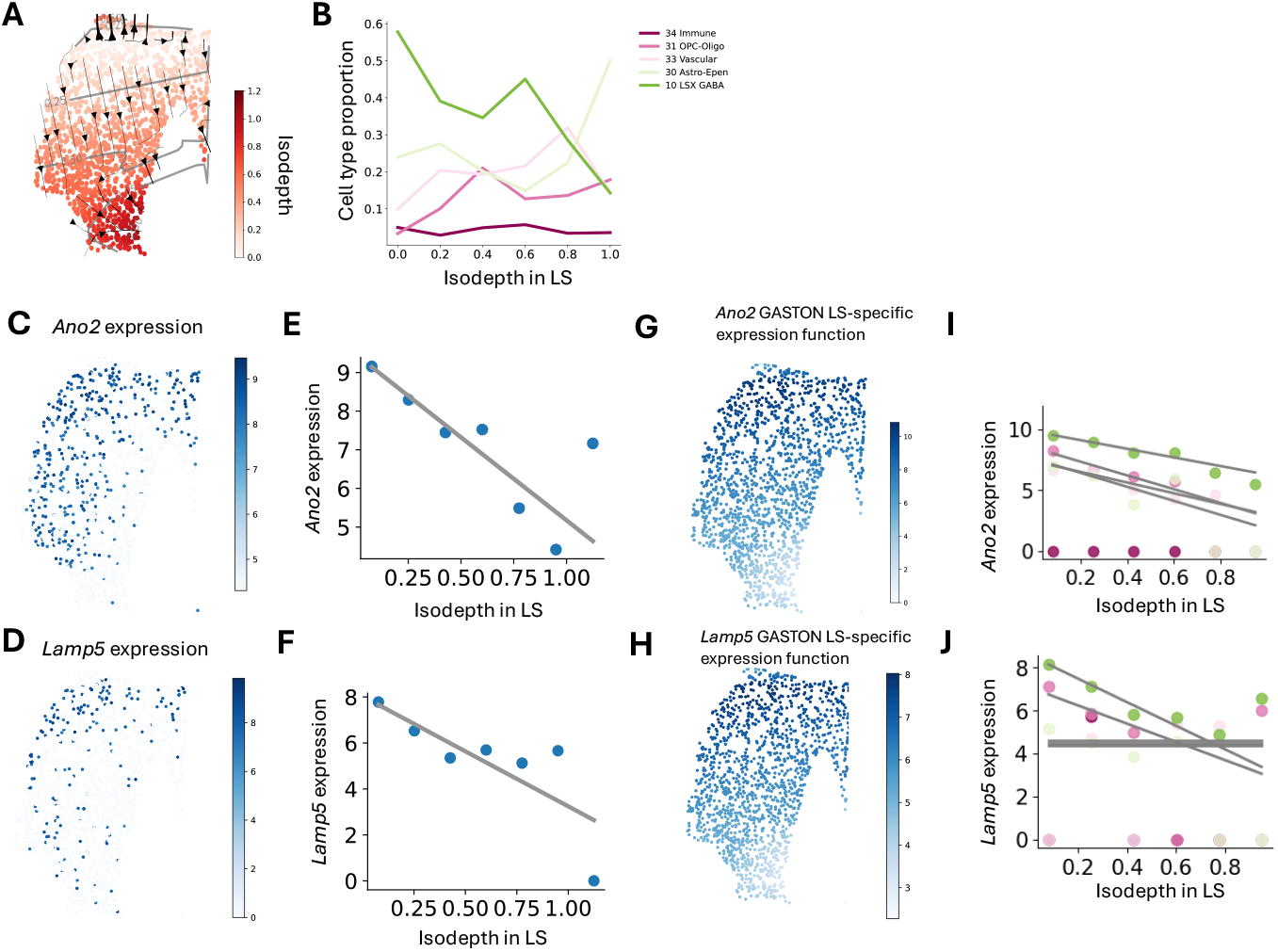
**(A)** Isodepth in lateral septum (LS) domain labeled in Figure 5B. **(B)** Cell type proportion versus GASTON-Mix-inferred isodepth in LS. **(C-J)** Same figures as in Figure S3 for genes *Ano2* and *Lamp5*.

**Figure S5:**
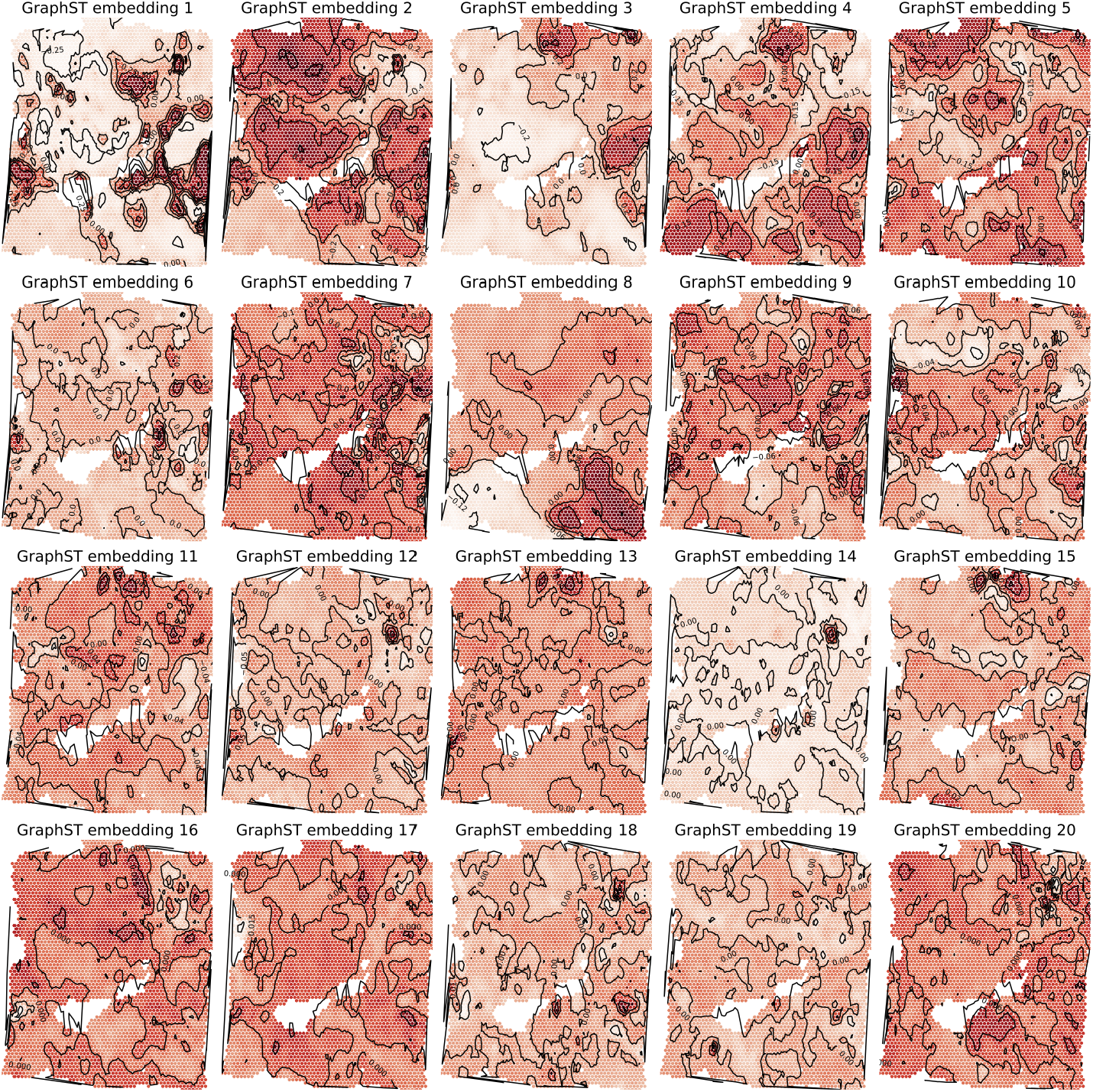
GraphST embedding components for 10x Visium breast cancer sample. Curves connect spots of equal embedding value.

